# Predicting genome sizes and restriction enzyme recognition-sequence probabilities across the eukaryotic tree of life

**DOI:** 10.1101/007781

**Authors:** Santiago Herrera, Paula H. Reyes-Herrera, Timothy M. Shank

## Abstract

High-throughput sequencing of reduced representation libraries obtained through digestion with restriction enzymes – generically known as restriction-site associated DNA sequencing (RAD-seq) – is a common strategy to generate genome-wide genotypic and sequence data from eukaryotes. A critical design element of any RAD-seq study is a knowledge of the approximate number of genetic markers that can be obtained for a taxon using different restriction enzymes, as this number determines the scope of a project, and ultimately defines its success. This number can only be directly determined if a reference genome sequence is available, or it can be estimated if the genome size and restriction recognition sequence probabilities are known. However, both scenarios are uncommon for non-model species. Here, we performed systematic *in silico* surveys of recognition sequences, for diverse and commonly used type II restriction enzymes across the eukaryotic tree of life. Our observations reveal that recognition-sequence frequencies for a given restriction enzyme are strikingly variable among broad eukaryotic taxonomic groups, being largely determined by phylogenetic relatedness. We demonstrate that genome sizes can be predicted from cleavage frequency data obtained with restriction enzymes targeting ‘neutral’ elements. Models based on genomic compositions are also effective tools to accurately calculate probabilities of recognition sequences across taxa, and can be applied to species for which reduced-representation data is available (including transcriptomes and ‘neutral’ RAD-seq datasets). The analytical pipeline developed in this study, PredRAD (https://github.com/phrh/PredRAD), and the resulting databases constitute valuable resources that will help guide the design of any study using RAD-seq or related methods.

## INTRODUCTION

The use of type II restriction enzymes to obtain reduced representation libraries from nuclear genomes, combined with the power of next-generation sequencing technologies, is rapidly becoming one of the most-used commonly strategies to generate genome-wide genotypic and sequence data in both model and non-model organisms (Baird et al. 2008; Andolfatto et al. 2011; Elshire et al. 2011; Peterson et al. 2012). The single nucleotide polymorphisms (SNPs) embedded in the resulting restriction-site associated DNA (RAD) sequence tags (M R Miller et al. 2007; Baird et al. 2008) have myriad uses in biology, which range from genetic mapping (Wang et al. 2013; Weber et al. 2013) to population genomics (Hohenlohe et al. 2010; Andersen et al. 2012; White et al. 2013), phylogeography (Emerson et al. 2010; Reitzel et al. 2013), phylogenetics (Wagner et al. 2012; Eaton & Ree 2013; Herrera et al. 2015; Herrera & Shank 2015), and SNP marker discovery (Scaglione et al. 2012; Toonen et al. 2013).

The choice of appropriate type II restriction enzyme(s) is critical for the effective design and application of RAD sequencing (RAD-seq) and a rapidly growing number of related methods such as genotyping-by-sequencing (Elshire et al. 2011), multiplexed shotgun genotyping (Andolfatto et al. 2011), double digest RAD-seq (Peterson et al. 2012), and ezRAD (Toonen et al. 2013). This choice determines the number of RAD markers that can be obtained, which in turn dictates the amount of sequencing needed for a desired coverage level, the number of samples that can be multiplexed, the monetary cost, and ultimately the success of a project. The theoretical maximum number of RAD markers that can be obtained for a given combination of restriction enzyme and biological species can be easily calculated as twice the frequency (absolute number of occurrences) of the enzyme’s recognition sequence (which for type II restriction enzymes is also the cleavage site) in the genome, but only when the fully sequenced genome is available. For cases in which the whole genome sequence is not available, i.e. most cases, this number can be approximated as twice the product of the genome size and the probability of the enzyme’s recognition sequence in a given genome.

### Genome sizes

Genome sizes can be approximated in non-model organisms through sequencing-independent techniques such as Feulgen densitometry (Hardie et al. 2002) or flow cytometry (Vinogradov 1994; Dolezel et al. 2007). However, these techniques have well known limitations: (1) flow cytometry often requires the availability of fresh tissue material with accessible intact cells or nuclei, thus diminishing its applicability to field-collected and fixed samples; and (2) both flow cytometry and Feulgen densitometry can be affected by staining interference with cytosolic compounds and variability in DNA packaging among cell types, which can significantly impact the accuracy and reproducibility of measurements (see reviews by Hardie *et al.* (2002) and Dolezel & Bartos (2005), and references therein). Therefore, alternative methods for genome-size estimation are desirable.

Type II restriction enzymes, which are endonucleases chiefly produced by prokaryotic microorganisms, cleave double stranded DNA (dsDNA) at specific unmethylated recognition sequences that are 4 to 8 base pairs long and typically palindromic. These enzymes are thought to play an important role as defense systems against foreign phage dsDNA during infection or as selfish parasitic elements, and therefore have been the center of an evolutionary ‘arms race’ (Rambach & Tiollais 1974; Rocha et al. 2001; Karlin et al. 1992). Type II restriction enzymes are not known in eukaryotes and are not used as virulence factors by bacteria to infect eukaryotic hosts. Therefore there are no *a priori* reasons to believe that recognition sequences in eukaryotic genomes are subject to selective pressures, but rather they should be evolutionarily neutral. A prediction from this neutrality hypothesis is that the frequency of restriction recognition sequences in a genome will be linearly correlated with the size of that genome, unless the particular restriction recognition-sequence is associated with non-neutral genomic elements. Hence, the genome size of a species can in theory be estimated from the number of markers obtained from a RADseq experiment, given that the restriction enzyme used shows the aforementioned linearity.

### Recognition-sequence probabilities

Flow cytometry has also been used as a sequencing-independent method to estimate the genomic guanine-cytosine (GC) composition (Vinogradov 1998; Šmarda et al. 2011), a widely suggested parameter for the estimation of the restriction enzyme’s recognition-sequence probability (Baird et al. 2008; Davey & Blaxter 2011). Nonetheless, preliminary evidence suggests that restriction recognition-sequence probability calculation, using GC composition as the only parameter, can yield predicted cleavage site frequencies that deviate significantly from observations, for particular combinations of taxa and restriction enzymes (Davey & Blaxter 2011; Davey et al. 2011). The extent and magnitude of these deviations across the eukaryotic tree of life remains unknown. Better models to calculate restriction recognition-sequence probabilities across taxonomic groups are needed to improve the accuracy of predictions of cleavage site frequencies in species without sequenced genomes. These models could be applied using non-genomic datasets (e.g., transcriptomes) to obtain recognition-sequence probability estimates, thus aiding the applicability of RAD-seq methods in non-model organisms.

Eukaryotic genomes have heterogeneous compositions with characteristic signatures at the level of di- and trinucleotides that are largely independent of coding status or function (Karlin & Mrázek 1997; Karlin et al. 1998; Gentles & Karlin 2001). Thus, it is possible that genome composition at these levels has a large influence on the abundance of short sequence patterns, such as recognition sequences of restriction enzymes. Models incorporating the information from these genomic compositional signatures should improve the accuracy of restriction recognition-sequence probability calculations.

Here we performed systematic *in silico* genome-wide surveys of genome compositions and recognition sequences, for diverse and commonly used type II restriction enzymes, in 434 eukaryotic whole and draft genomes (Supplementary Table 1) to: (1) characterize restriction recognition-sequence frequencies across the eukaryotic tree of life; (2) explore the potential for predicting genome sizes from restriction recognition-sequence frequency data; (3) develop stochastic models based on genomic compositions to calculate probabilities of recognition sequences across taxa; and (4) evaluate the applicability of these models to species for which only non-genomic data is available (i.e., not whole or draft genome assemblies), such as transcriptomes or restriction-site associated DNA sequence data. The PredRAD analytical pipeline developed in this study (https://github.com/phrh/PredRAD), and the resulting databases constitute a valuable reference resource that will help guide restriction enzyme choice infuture studies using broadly applicable RAD-related methods.

**Table 1.**
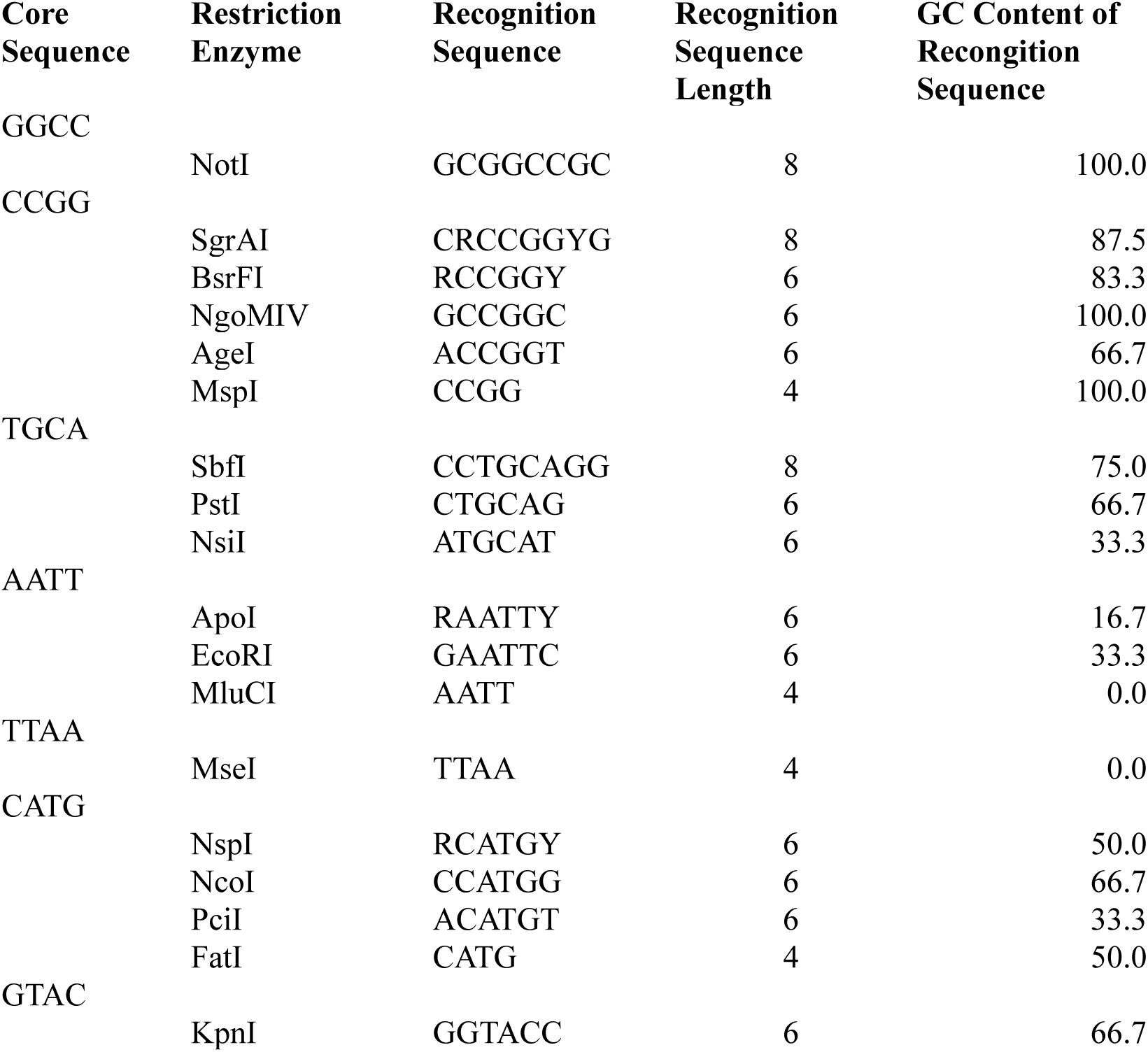
Restriction enzymes included in this study.

## RESULTS

### Frequencies of recognition sequences are highly variable across taxa

To characterize cleavage site frequencies across the Eukaryotic tree of life, we surveyed restriction recognition-sequences for 18 commonly used palindromic type II restriction enzymes in 434 whole and draft genomes. Observed relative frequencies of recognition sequences were highly variable among broad taxonomic groups for the set of restriction enzymes here examined (Table 1) – except for FatI – with clear clustering patterns determined by phylogeny (Figure 1). For example, with NgoMIV we observed 45.8 recognition sequences per megabase (RS/Mb) ± 24.6 (mean ± SD) in core eudicot plants, compared to 277.4 ± 131.3 RS/Mb in commelinid plants (monocots). Among closely-related species the relative frequency patterns were similar and variability generally small. Observed relative frequencies of RS/Mb were inversely proportional to the length of the recognition sequence, with orders of magnitude differences among the 4-, 6-, and 8-cutters when compared within the same species, e.g., in the starlet anemone *Nematostella vectensis* there were 3917.6, 167.6, and 6.9 RS/Mb for the 4-cutter FatI, 6-cutter PstI and 8-cutter SbfI, respectively. In contrast, nucleotide composition of the recognition sequence itself did not show a clear correlation with the observed relative frequency of cleavage sites. For example, 83.6 RS/Mb ± 25.1 were observed in Neopterigii vertebrates for KpnI (GGTACC) and 622.6 RS/Mb ± 119.1 were observed for PstI (CTGCAG), both recognition sequences with the same GC content (66.7%).

**Fig. 1.**
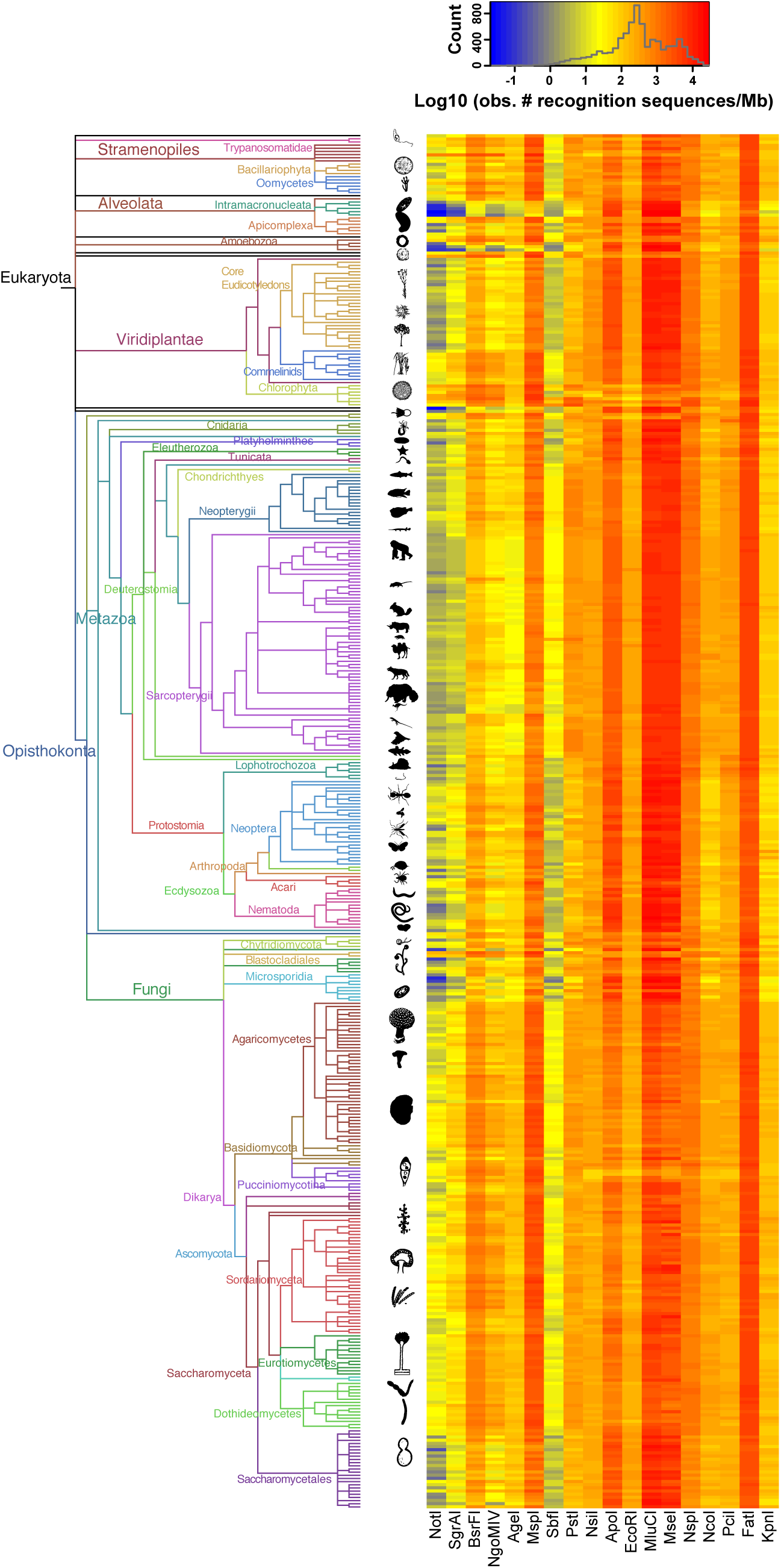
Observed restriction recognition-sequence frequencies. Left: phylogenetic tree of all eukaryotic taxa analyzed in this study. The tree is based on the NCBI taxonomy tree retrieved on May 16, 2013 using the iTOL tool http://itol.embl.de. Branch colors and labels indicate broad taxonomic groups. Organism silhouettes and cartoons were created by the authors or obtained from http://phylopic.org/. Right: heatmap of the observed frequency of restriction sites. Each row corresponds to a species from the tree on the left, and each column corresponds to a different restriction enzyme. Gray line in the color-scale box shows the distribution histogram of all values.

### Genome sizes can be predicted from particular recognition-sequence frequencies

To explore the potential for predicting genome sizes from restriction recognition-sequence frequency data, we modeled their relationship using data from the 434 genomes and 18 restriction enzymes through linear regression. A general positive correlation between recognition-sequence frequency and genome size was observed for all restriction enzymes, being significantly strong (Spearman’s correlation coefficient >0.95) for five of them: EcoRI, FatI, NsiI, NspI and PciI (Figure 2). Predicted genome sizes, calculated using the linear models with estimated beta parameters for these five enzymes (Table 2), matched actual observed genome size values extremely well (Supplementary Figure 1).

**Table 2.** Linear regresion parameter estimates and 95% confidence intervals.

| **Enzyme** | **B0** | **B1** | **95% CI B0** | **95% CI B1** |
| --- | --- | --- | --- | --- |
| AgeI | 3.791226 | 1.081123 | (3.589084,3.993368) | (1.030957,1.131288) |
| ApoI | 3.909432 | 0.789828 | (3.771043,4.047820) | (0.764083,0.815571) |
| BsrFI | 3.595150 | 0.972785 | (3.336245,3.854053) | (0.917377,1.028193) |
| EcoRI | 3.915725 | 0.932289 | (3.836952,3.994497) | (0.914985,0.949591) |
| FatI | 2.719837 | 0.947207 | (2.639872,2.799802) | (0.933248,0.961165) |
| KpnI | 4.041810 | 0.984192 | (3.931500,4.152119) | (0.957826,1.010558) |
| MluCI | 3.432945 | 0.796619 | (3.281188,3.584701) | (0.770985,0.822252) |
| MseI | 3.963499 | 0.722786 | (3.813020,4.113977) | (0.696835,0.748737) |
| MspI | 3.084383 | 0.957434 | (2.846370,3.322395) | (0.912357,1.002510) |
| NcoI | 4.089533 | 0.910311 | (3.975127,4.203937) | (0.884724,0.935898) |
| NgoMIV | 5.115077 | 0.738618 | (4.881512,5.348642) | (0.681804,0.795430) |
| NotI | 6.432067 | 0.581703 | (6.254678,6.609455) | (0.522412,0.640993) |
| NsiI | 3.948432 | 0.908376 | (3.874564,4.022299) | (0.892446,0.924304) |
| NspI | 3.399772 | 0.930233 | (3.316885,3.482657) | (0.914012,0.946453) |
| PciI | 4.092091 | 0.885098 | (4.031567,4.152614) | (0.871942,0.898254) |
| PstI | 4.244698 | 0.850488 | (4.114538,4.374857) | (0.822215,0.878759) |
| SbfI | 5.782031 | 0.726905 | (5.671977,5.892083) | (0.693729,0.760080) |
| SgrAI | 5.500710 | 0.749462 | (5.245991,5.755428) | (0.677348,0.821576) |

**Fig. 2.**
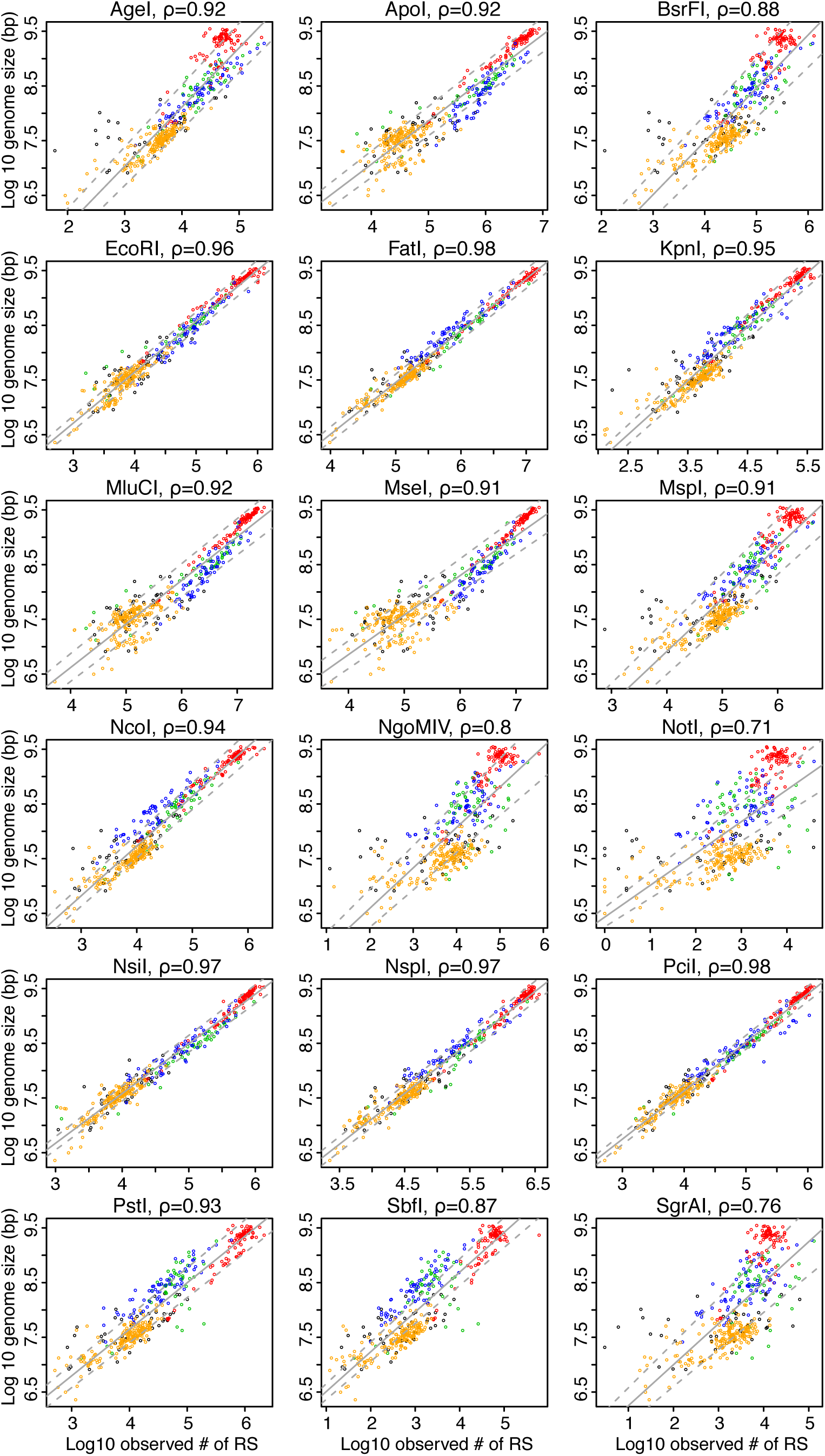
Linear correlations of restriction recognition-sequence frequencies and genome sizes. Scatter plots show the observed numbers of recognition sequences in a genome for a given restriction enzyme (x-axis) vs. the size of the genome in base pairs (y-axis). The data for all the 434 examined genomes are shown. Each panel shows the data for a different restriction enzyme. Dot colors indicate broad taxonomic groups: fungi (yellow), plants (green), invertebrates (blue), vertebrates (red), and others (black). Non-parametric Spearman’s rank-order correlation coefficients (*ρ*) are shown for each restriction enzyme. Solid gray lines represent the best-fit linear models with 95% confidence intervals (gray dotted lines).

### Genome composition-based models outperform traditional GC content-based models

To generate better models for cleavage site frequency calculation in species without sequenced genomes, we developed stochastic models based on the GC content of each genome, as well as the mononucleotide, dinucleotide and trinucleotide compositions to predict the expected frequency of recognition sequences for each restriction enzyme. We evaluated the fit of each model by comparing the *in silico* observed frequencies of cleavage sites to the expected frequencies predicted by the models using a similarity index (*SI),* defined as the quotient of the number of observed and expected cleavage sites, minus one. A positive *SI* indicates that the number of observed cleavage sites is greater than the expected, whereas a negative *SI* indicates a smaller number of observed sites than expected. If *SI* is equal to 0, then the number of observed sites is equal to the expectation. For example, a *SI* = 1 indicates that the number of observed cleavage sites for a particular enzyme in a given genome is twice the number of expected sites predicted by a particular model. Trinucleotide composition models were in general a better predictor, in terms of their accuracy and precision, of the expected number of cleavage sites than any of the other models (Figures 3, 4 and 5). The mononucleotide and GC content models produced relatively poorer predictions that were indistinguishable from one another (Figures 3, 4 and 5). In a few cases the other models outperformed the trinucleotide model, e.g., EcoRI (Figures 3, 4 and 5). The fit of the predictions was highly variable among broad taxonomic groups but generally similar within, e.g., in Neopterigii vertebrates an average *SI* of 0.14 ± 0.19 for AgeI with the dinucleotide model, compared to -0.31 ± 0.19 in Sarcopterigii.

**Fig. 3.**
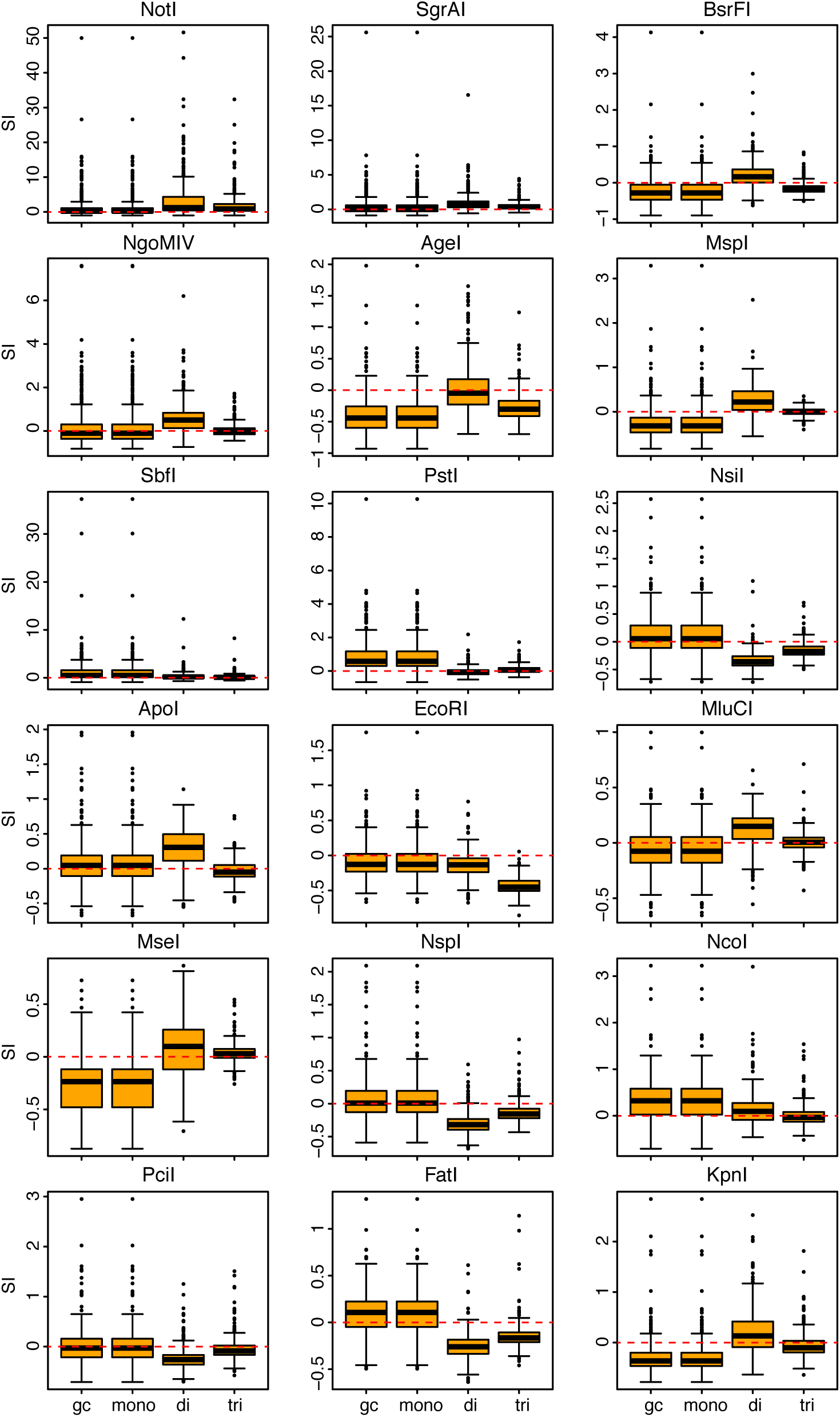
Overall fit of genome composition models per restriction enzyme. Vertical axes in the box and whisker plots indicate the values of the similarity index (*SI*) for each species per enzyme (see Methods section). Horizontal axes in the box and whisker plots indicate the genome composition model: GC content (gc), mononucleotide (mono), dinucleotide (di), and trinucleotide (tri). Horizontal edges of range boxes indicate the first and third quartiles of the *SI* values under each composition model. The thick horizontal black line represents the median. Whiskers indicate the value of 1.5 times the inter-quartile range from the first and third quartiles. Outliers are defined as SI values outside the whiskers range and are represented by dots. Outlier value of *Entamoeba histoyitica* for *NotI* was excluded. Red dotted lines indicate *SI*=0.

**Fig. 4.**
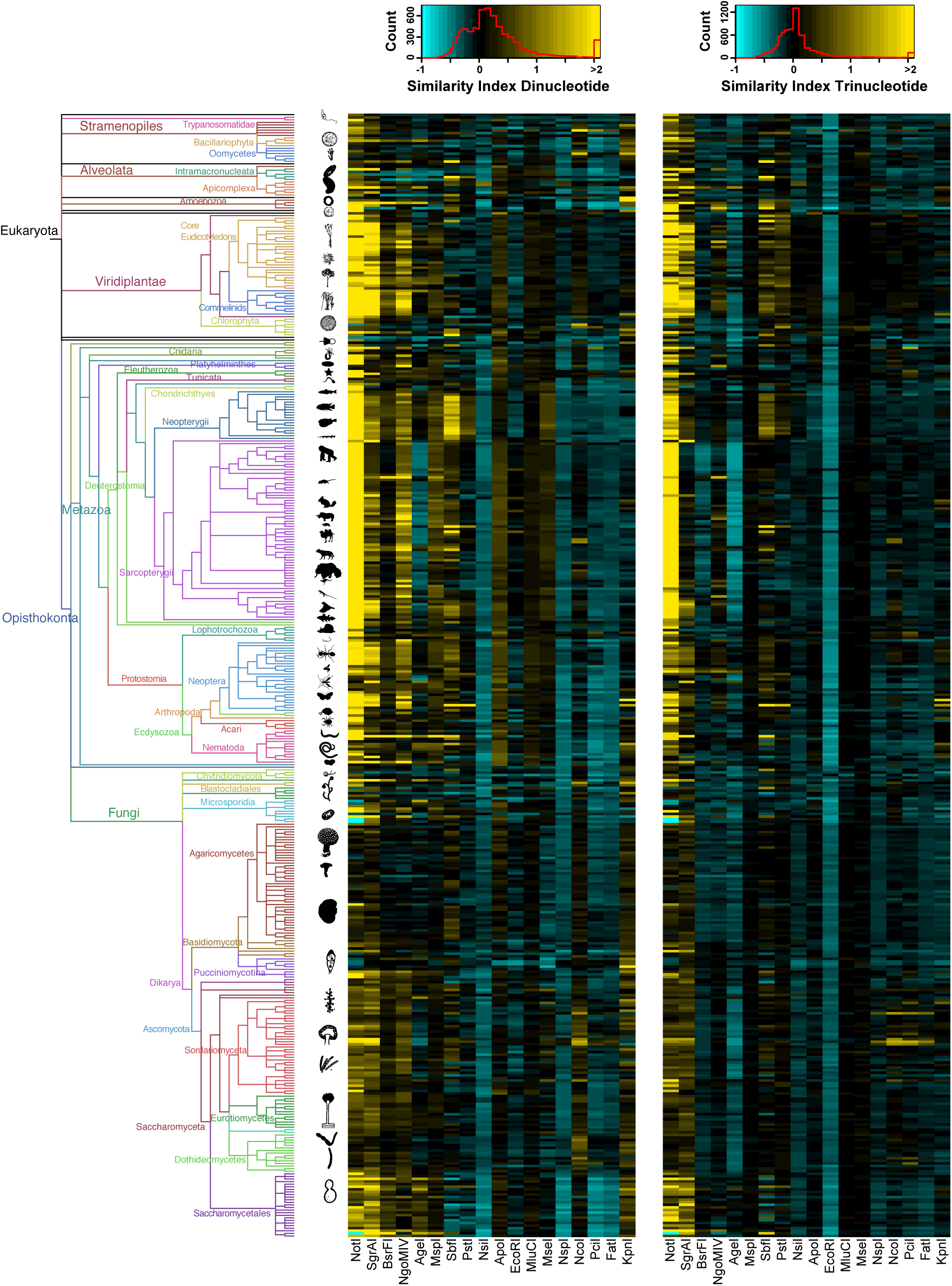
Similarity indexes for dinucleotide and trinucleotide genome composition models. Left: phylogenetic tree as in Fig 1. Center: heatmap of the similarity indexes for the dinucleotide model Right: heatmap of the similarity indexes for the mononucleotide model. Each row corresponds to a species from the tree on the left, and each column corresponds to a different restriction enzyme. Cyan indicates *SI <* 0 and yellow indicates *SI >* 0. Red line in the color-scale box shows the distribution histogram of all values.

**Fig. 5.**
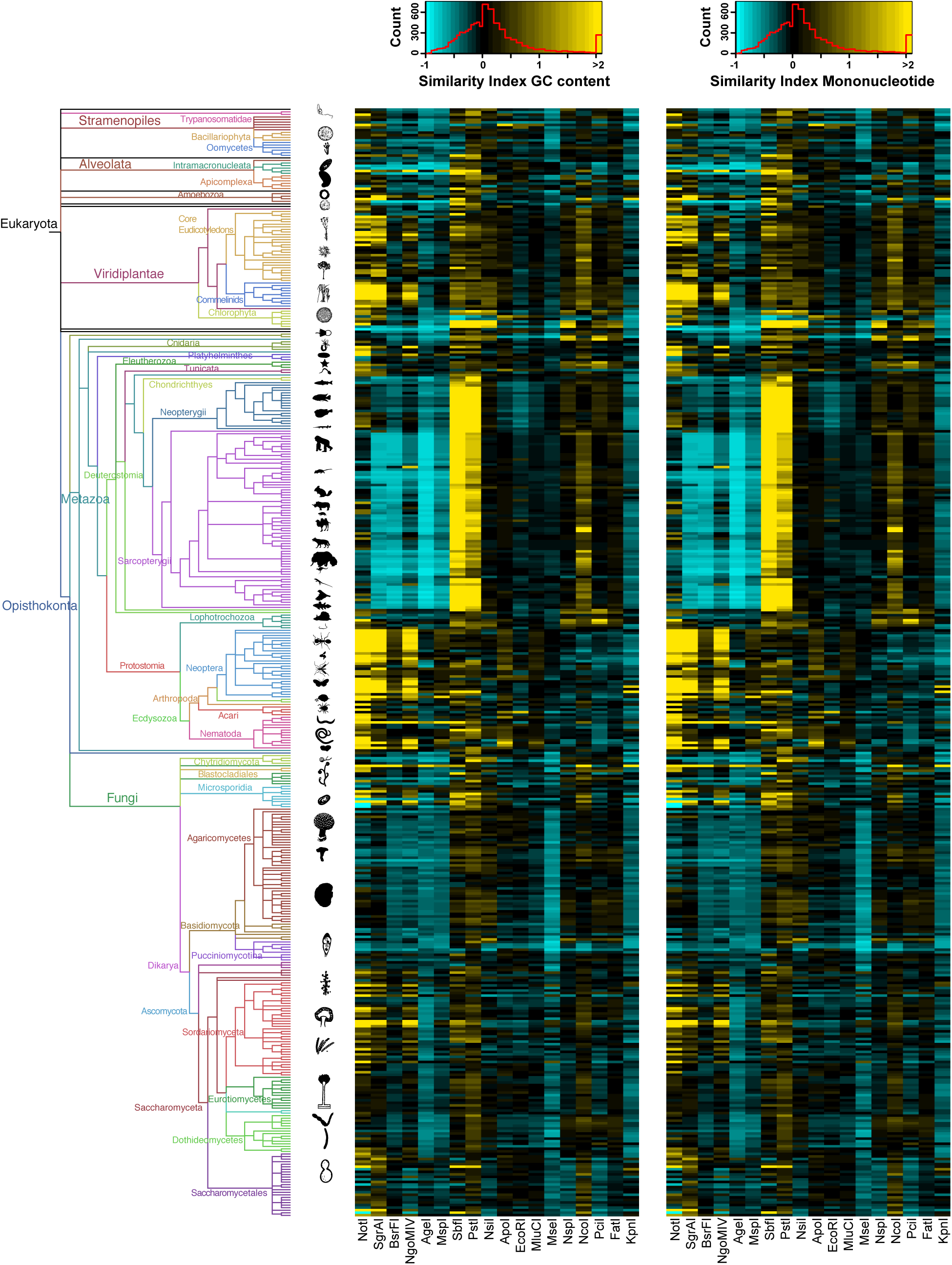
Similarity indexes for GC content and mononucleotide genome composition models. Left: phylogenetic tree as in Fig 1. Center: heatmap of the similarity indexes for the GC content model Right: heatmap of the similarity indexes for the mononucleotide model. Each row corresponds to a species from the tree on the left, and each column corresponds to a different restriction enzyme. Cyan indicates *SI <* 0 and yellow indicates *SI >* 0. Red line in the color-scale box shows the distribution histogram of all values.

### Recognition-sequence probability can be calculated from non-genomic datasets

Genomic resources (whole or draft genomes) are unavailable for most species (Dunn & Ryan [https://bitbucket.org/caseywdunn/animal-genomes] estimate that ∼0.015% of species have a published genome to date). However, reduced representation datasets that capture a small fraction of a genome, such as RNA-seq or RAD-seq datasets, can now be easily and economically developed. We investigated the potential use of these datasets to estimate genome composition parameters for our predictive models and calculate recognition-sequence probabilities of any given restriction enzyme. For this we selected a set of 27 species out of the 434 examined eukaryotic species with whole and draft genomes, which also have publically available transcriptome data. The restriction-sequence probabilities calculated for the same panel of 18 restriction enzymes, as above, were remarkably similar between those calculated using known composition parameters from the whole and draft genomes and those calculated using estimated composition parameters from transcriptome datasets (Figure 6). Interestingly, the overall similarity between the two kinds of calculated probabilities (measured as the mean squared error – MSE – calculated across all species) was greatest when probabilities were calculated using a mononucleotide composition model (0.046; when MSE=0 the probabilities are identical; MSE value increases as similarity decreases), and decreased when dinucleotide and trinucleotide models were used (0.06 and 0.07, respectively). As expected, the species’ specific MSE values were variable, and tended to decrease as the propotion of genome represented by the transcriptome increased (Figure 6).

**Fig. 6.**
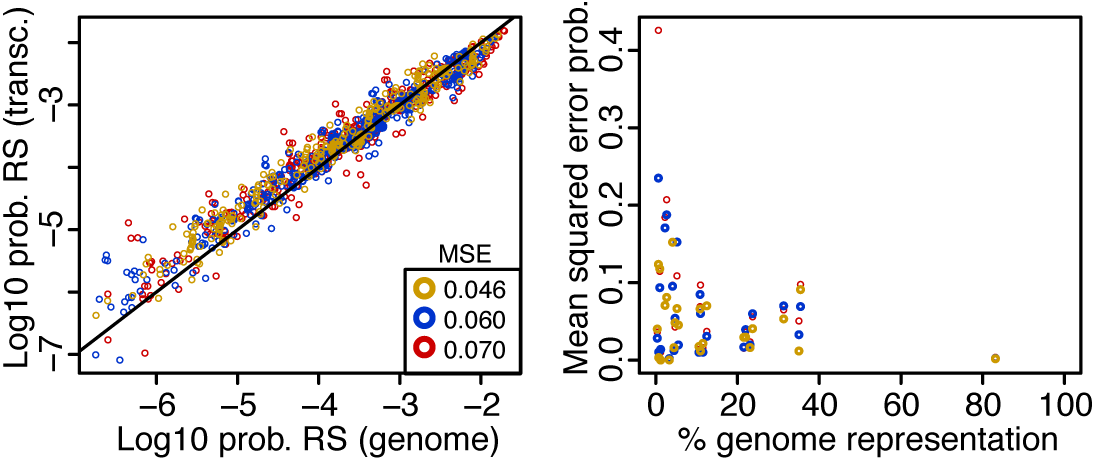
Left: Scatter plot of the probability of restriction recognition-sequence (RS) probabilities calculated using known composition parameters from the genome (x-axis) vs. those calculated using estimated composition parameters from transcriptome datasets (y-axis). Each dot represents the combination of one of the 18 examined restriction enzymes and one of the 27 species in the reduced-representation subset. Colors indicate the probabilities calculated by different models: mononucleotides (yellow), dinucleotides (blue), trinucleotides (red). Average mean squared error (MSE) values for the probabilities calculated with each model are shown. Solid black line represents the identity line, in which x = y. Right: Scatter plot of the percentage of the genome represented by the transcriptome datasets (x-axis) vs. per-species mean squared error (MSE) values for the probabilities calculated with each model (y-axis). As before, colors indicate the probabilities calculated by different models: mononucleotides (yellow), dinucleotides (blue), trinucleotides (red).

We also calculated recognition-sequence probabilities using parameters estimated from the *in silico* RAD sequence data for the same 27 species, finding great variability (Figure 7). The recognition-sequence probabilities calculated using parameters estimated from RAD-seq datasets obtained with enzymes that showed strong correlations between recognition-sequence frequency and genome size (Figure 2) were almost identical to the probabilities calculated using the known composition parameters from the whole or draft genome datasets (Figure 7, Supplementary Figure 2). Contrastingly, the probabilities calculated from RAD-seq datasets obtained with enzymes that showed weaker correlations between recognition-sequence frequency and genome size (such as NotI, NgoMIV and SgrAI) were substantially dissimilar (Figure 7, Supplementary Figure 2). Overall, as observed for the transcriptome datasets, the similarity between the two kinds of calculated probabilities (measured by the MSE) was greatest when probabilities were calculated using a mononucleotide composition model, and decreased when dinucleotide and trinucleotide models were used. Similarly, the species-specific MSE values tended to decrease as the proportion of genome represented by the RAD-seq datasets increased (Supplementary Figure 2), although in some cases (e.g. PstI and SbfI) they showed a marked decrease followed by an increase at higher representation proportions.

**Fig. 7.**
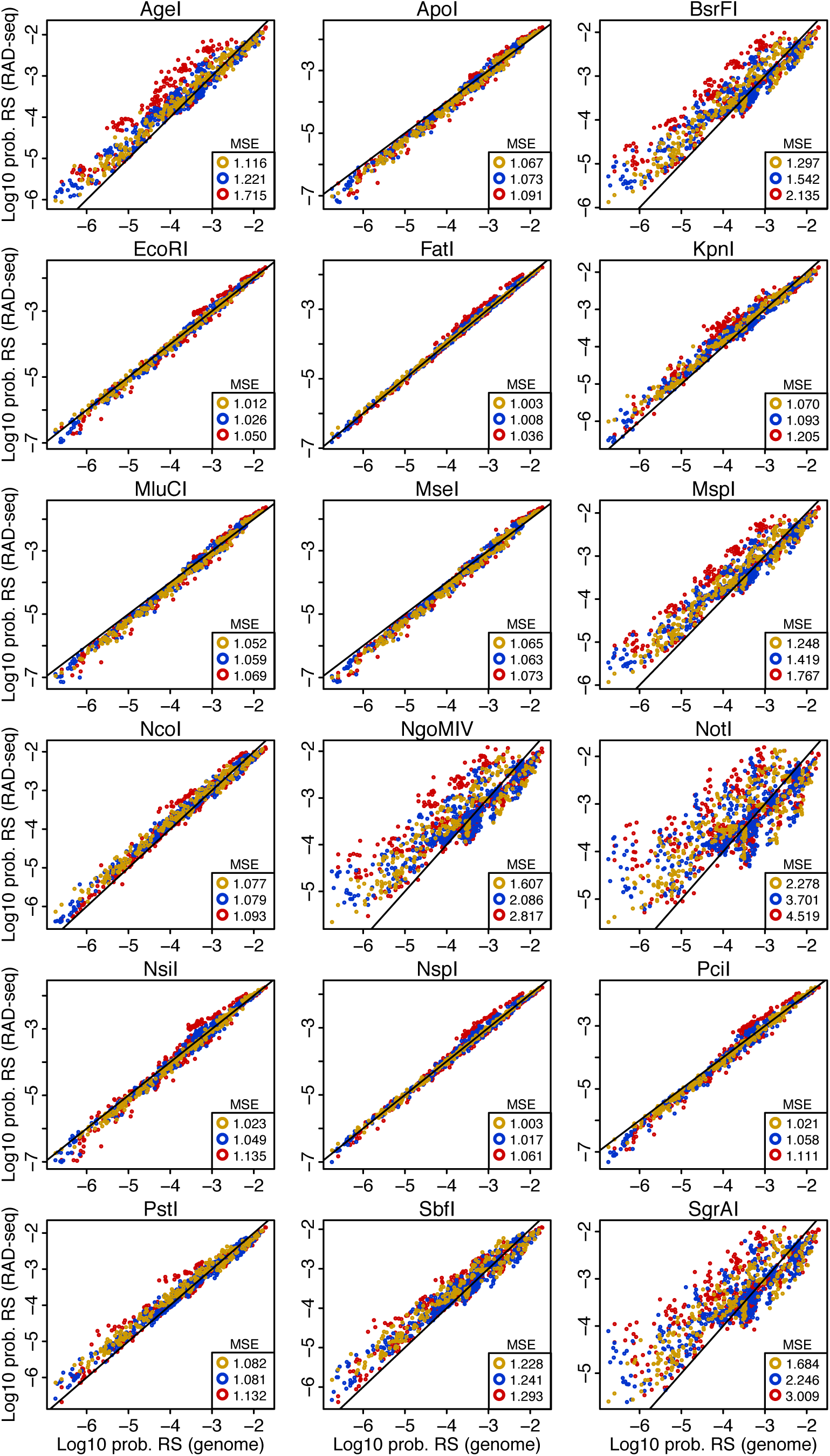
Scatter plots of the probability of restriction recognition-sequence (RS) probabilities calculated using known composition parameters from the genome (x-axis) vs. those calculated using estimated composition parameters from *in silico* RAD-seq datasets (y-axis). Each dot represents the combination of one of the 18 examined restriction enzymes and one of the 27 species in the reduced-representation subset. Colors indicate the probabilities calculated by different models: mononucleotides (yellow), dinucleotides (blue), trinucleotides (red). Average mean squared error (MSE) values for the probabilities calculated with each model are shown. Solid black lines represents the identity lines, in which x = y.

Predicted frequencies of cleavage sites (absolute number of cleavage sites), obtained by multiplying known genome sizes with the probabilities calculated using composition models, were remarkably similar to the observed frequencies of cleavage sites in whole and draft genome datasets when the model parameters were estimated from transcriptome datasets, or from RAD-seq datasets generated with restriction enzymes showing strong correlations between recognition-sequence frequency and genome size (Supplementary Figures 3, 4 and 5).

## DISCUSSION

### Genome-wide surveys of cleavage sites across the eukaryotic tree of life

Observed recognition-sequence frequencies for a given restriction enzyme are strikingly variable across broad eukaryotic taxonomic groups, but are similar among closely-related species. This finding is consistent with the hypothesis that the abundance of cleavage sites is largely determined by phylogenetic relatedness. This pattern is most evident in groups that have a larger taxonomic representation, such as mammals. As more genome assemblies become available, patterns within many other underrepresented taxonomic groups will be clarified. Through the use of comparative methods in a robust phylogenetic framework, it will be possible to establish taxon-specific divergence thresholds that could diagnostic of significant evolutionary changes in genome architecture.

As expected, observed relative frequencies of cleavage sites with shorter recognition sequences are on average higher than the observed frequencies with longer recognition sequences. However, this pattern in not universal. There are several instances in which the relative frequency of cleavage sites for a high-denomination cutter is higher than that for a low-denomination cutter. For example, in primates the relative frequency of the 8-cutter SbfI (24.6 ± 1.7 RS/Mb) is significantly higher than the frequency of the 6-cutter AgeI (18.4 ± 1.4 RS/Mb). These deviations from expectation are indicative of enzyme-specific frequency biases for particular taxa, and, as illustrated in the results section, are not correlated with the base composition of recognition sequences. These observations demonstrate that the expected relative frequencies of recognition sequences cannot be naively extrapolated across enzyme types and divergent taxa, but rather, specific knowledge of recognition sequence frequencies/probabilities and genome sizes is needed.

### Predictability of genome sizes

For many of the examined type II restriction enzymes examined (e.g. EcoRI, FatI, NsilI, NspI, PciI), the observed frequencies of recognition sequences in eukaryotic genomes are consistent with the idea that they behave neutrally, evolutionary speaking, and therefore can be readily used as parameters in linear models to estimate genome sizes (Figures 1 and 2). In contrast, the observed frequencies of recognition sequences for some other type II restriction enzymes showed significant deviations from the predictions of this evolutionary neutrality hypothesis (e.g. BsrFI, NgoMIV, NotI, SbfI, SgrAI). A closer look at the genomic locations of the recognition sequences of these deviant cases reveals that, in mammals, they are more likely to occur in conserved-element genomic regions than what would be expected by chance (Figure 8). Conserved genomic elements (*sensu* Siepel *et al.* 2005) are widely recognized as evidence of functional regions, mainly regulatory, under strong purifying (negative) selection (Katzman et al. 2007; Bejerano et al. 2004). Thus, this observation suggests that the association of some restriction recognition sequences with non-neutral genomic elements in particular taxa can account for some of the observed biases and heterogeneity in the relative frequencies of cleavage site across the eukaryotic tree of life. Further comparative genomic studies in underrepresented clades promise to unravel additional potential mechanisms that can further explain observed deviations from expected neutral behavior.

**Fig. 8.**
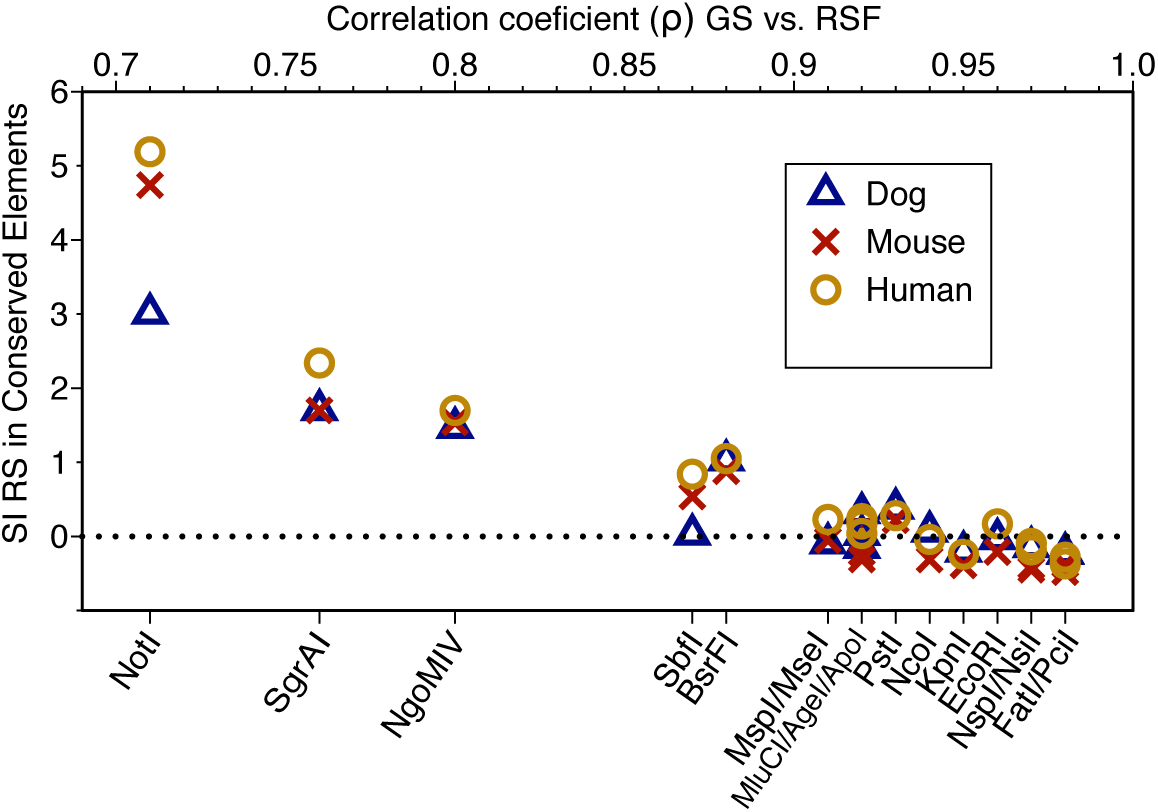
Scatter plot of the Spearman’s correlation coefficient (*ρ*) values between genome sizes and restriction recognition-sequence frequencies (x-axis) vs. similarity index (SI) values of between observed and expected numbers of restriction recognition-sequences in mammalian conserved-element genomic regions (y-axis). Corresponding restriction enzyme names to each *ρ* value are shown. Colors and symbols indicate different taxa: dog (blue triangles), mouse (red crosses), human (yellow circles). Dotted line indicates SI = 0 (value at which expected and observed values are equal).

### Predictability of recognition sequence probabilites

Our analyses indicate that in most cases stochastic models based on trinucleotide compositions are the best predictors, and the GC content and mononucleotide models are the worst predictors of the expected relative number of cleavage sites in a eukaryotic genome. It is likely that the greater number of parameters in the trinucleotide model (64, compared to 16, 4 and 2 of the dinucleotide, mononucleotide and GC content model, respectively), combined with the greater k-mer length, is the cause of the better fit. However, this trend is not universal. As illustrated in the results section, in a few cases the other models outperformed the trinucleotide composition model. Neither the GC content nor length of the recognition sequence can confidently explain the observed discrepancies. Increasing the k-mer length above trinucleotide in the composition models (i.e., tetranucleotide, pentanucleotide, etc.) could improve their fit, however this will come at a cost of increasing probability calculation error in reduced-representation datasets (caused by sampling error in datasets composed by many short contigs) (Figures 6 and 7, Supplementary Figure 2). Future cost-benefit evaluations of the overall influence of these factors (k-mer length and genome sampling error) will help elucidate their relative contributions to recognition sequence probability calculations using parameters estimated from non-genomic reduced-representation datasets.

It is not surprising that the fit of the predictions made by the models is highly variable across taxonomic groups, given the high heterogeneity observed in the genetic composition patterns across the eukaryotic tree of life (Appendix I). We conclude that the predictability of cleavage site frequencies in eukaryotic genomes needs to be treated on a case-specific basis, whereby the phylogenetic position of the taxon of interest, its genome size, and the probability of the recognition sequence of the selected restriction enzyme are the chief foci among the most determinant factors.

The remarkable similarity between probabilities calculated using parameters estimated from nongenomic (transcriptome and ‘neutral’ RAD-seq datasets) and genomic datasets demonstrates the potential of using extant reduced-representation datasets for planning further restriction site associated DNA sequencing projects. Although transcriptome datasets by definition are enriched in functional genomic regions (transcribed genes) that are known to be targets of natural selection at different levels (codons, protein domains, etc.) we find no evidence of substantial differences in the underlying mononucleotide, dinucleotide and trinucleotide compositions compared to the overall genome-wide compositions. This observation is consistent with previous studies showing that genomic composition does not vary significantly between non-coding and coding regions (Karlin & Mrázek 1997; Karlin et al. 1998; Gentles & Karlin 2001). In the cases of RAD-seq datasets, there are a clear biases in the underlying mononucleotide, dinucleotide and trinucleotide compositions for datasets generated with restriction enzymes targeting ‘non-neutral’ recognition sequences (e.g., NotI, SgrAI, NgoMIV, SbfI, BsrFI) (Figures 7 and 8), compared to the overall genome-wide compositions, as evidenced by the calculated recognition sequence probabilities. As discussed above, these biases are likely caused in part by associations with conserved regions under strong selective pressures. RAD-seq datasets generated with restriction enzymes that are known to target ‘non-neutral’ recognition sequences should not be utilized for genome size estimation and restriction recognition-sequence probability calculation as these would likely yield biased inferences.

### Applications to study design with RAD-seq and related methodologies

For the design of a study using RAD-seq, or a related methodology, there are two fundamental questions that researchers commonly face: (1) what is the best restriction enzyme to use to obtain a desired number of RAD tags in the organism of interest? and (2) how many markers can be obtained with a particular enzyme in the organism of interest? The results from this study coupled with the developed software pipeline PredRAD, will allow any researcher to obtain an approximate answer to these questions. The flow diagram in Figure 9 illustrates a suggested workflow.

**Fig. 9.**
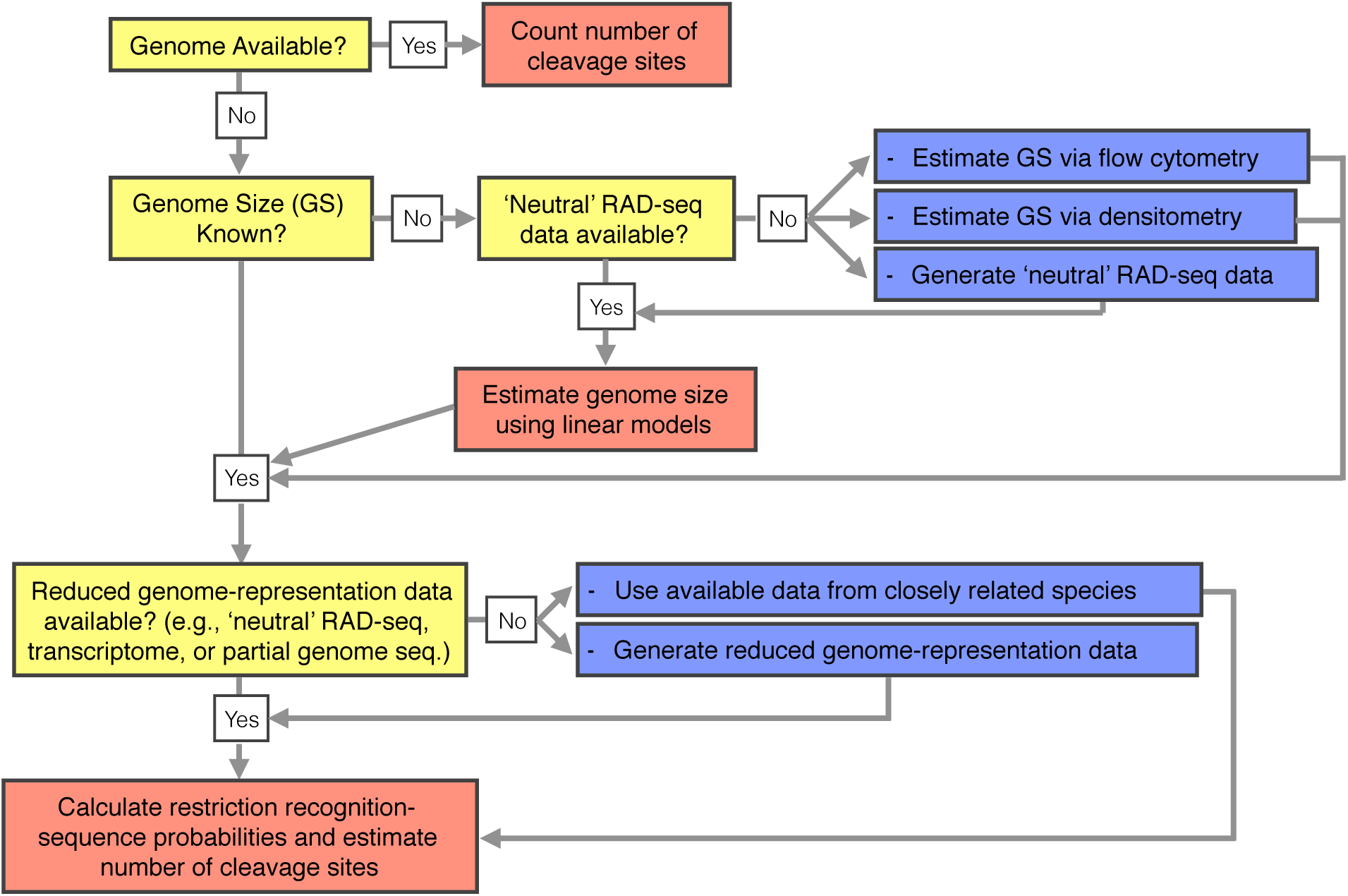
Suggested workflow to obtain an approximation to the number of cleavage sites for a set of restriction enzymes in a species of interest. Yellow boxes indicate question checkpoints, blue boxes indicate experimental steps, and red boxes indicate computational steps that can be carried out with the PredRAD analytical pipeline.

In a best-case scenario for the practical design of a study using RAD-seq, or a related methodology, the species of interest is already included in the database presented here. In this case, the best proxy for the estimated number of RAD tags that could be obtained empirically would be twice the number of *in silico* observed cleavage sites for each restriction enzyme (each cleavage site is expected to produce two RAD tags, one in each direction from the cleavage site) minus the number of *in silico* tags that align to multiple regions in the genome. For most of the 434 genomes examined in this study the recovery of RAD-tags after *in silico* sequencing was notably high, with a median percentage of suppressed alignments to the reference genome assembly of only 3% (Supplementary Figure 6). We observed no evident recovery bias by restriction enzyme, but rather bias was pronounced in a few individual species, likely indicating an enrichment of repetitive regions or duplications. For library preparation protocols in which a fragment size selection step is done without a prior shearing step, e.g., ddRAD (Peterson et al. 2012) and ezRAD (Toonen et al. 2013), the *size.select* function of the software package SimRAD (Lepais & Weir 2014) constitutes a valuable complementary study-design tool. If a new genome assembly becomes available for the target species and/or the researcher wishes to evaluate an additional restriction enzyme, PredRAD can be utilized with these data to quantify the number of cleavage sites and the recovery potential, as well as to estimate the probability of the new recognition sequence based on genome composition models.

In the scenario that the genome sequence of the species of interest is not available, the genome size and restriction recognition-sequence probability for the enzyme(s) of interest can be estimated to obtain an approximation of recognition-sequence frequencies (absolute numbers). Our observations demonstrate that a genome-size range can be estimated by applying linear regression models to the number of markers obtained in an empirical RAD-seq experiment using a restriction enzyme targeting a ‘neutral’ recognition sequence (e.g., EcoRI, NsilI, NspI, PciI; we advise caution using 4-cutter enzymes as in some taxa they can have cleavage frequencies that may effectively lead to sequencing the whole genome through RAD-seq [i.e., more than one cleavage site per 100bp]). Alternatively, genome size can also be estimated via flow cytometry and/or Feulgen densitometry (Vinogradov 1994; Dolezel et al. 2007; Hardie et al. 2002) for comparison. A range of restriction recognition-sequence probabilities can be obtained through genome composition models using parameters estimated from non-genomic reduced-representation datasets, such as transcriptomes, ‘neutral’ RAD-seq datasets, or even partial genome sequences, for the species of interest. Non-genomic datasets from closely related species could also be used to estimate these parameters, although the effect of evolutionary divergence on compositional differences warrants further exploration. Similarly, examination of other restriction enzymes with diverse recognition sequences, in addition to the ones examined in this study, promises great potential to identify ‘gold standard’ sets of enzymes for groups of taxa, with the goal of obtaining ‘neutral’ RAD-seq datasets.

Although genome size and the relative frequency (probability) of restriction recognition sequences are arguably the main determinant factors influencing the number of RAD tag markers that can be obtained experimentally, there are other factors that need to be considered during study design and data analysis steps. These include: genome differences among individuals, level of heterozygosity, the amount of methylation and other DNA modifications in the genome, the sensitivity of a particular cleavage enzyme to methylation and other DNA modifications, the efficiency of the enzymatic digestion, the number of repetitive regions and gene duplicates present in the target genome, the quality of library preparation and sequencing, the amount of sequencing, sequencing and library preparation biases, and the parameters used to clean, cluster and analyze the data, among others (see Davey *et al.* (2013), Catchen *et al.* (2013), DaCosta & Sorenson (2014), and Mastretta-Yanes *et al.* (2014) for further discussions).

## CONCLUSIONS

In this study, we performed systematic *in silico* genome-wide surveys of genome compositions and recognition sequences, for diverse and commonly used type II restriction enzymes across the eukaryotic tree of life. Our observations reveal that recognition-sequence frequencies for a given restriction enzyme are strikingly variable among broad eukaryotic taxonomic groups, being largely determined by phylogenetic relatedness. We demonstrate that genome sizes can be predicted from cleavage frequency data obtained with restriction enzymes targeting ‘neutral’ recognition sequences. Stochastic models based on genomic compositions are also effective tools to accurately calculate probabilities of recognition sequences across taxa, and can be applied to species for which reduced-representation genomic data is available (including transcriptomes and ‘neutral’ RAD-seq datasets). The results from this study and the software developed from it will help guide the design of any study using RAD sequencing and related methods. As more genome assemblies become available in underrepresented taxonomic groups, the patterns of compositional biases and restriction site frequencies across the eukaryotic tree of life will become clearer and will improve our understanding of genome evolution.

## MATERIALS & METHODS

### Observed restriction recognition-sequence frequencies

Assemblies from eukaryotic whole genome shotgun (WGS) sequencing projects available as of December 2012 were retrieved primarily from the U.S. National Center for Biotechnology Information (NCBI) WGS database (Supplementary Table 1). Only one species per genus was included. Of the 434 genome assemblies included in this study, 42% corresponded to fungi, 21% to vertebrates, 16% invertebrates, and 9% plants. Only unambiguous nucleotide calls were taken into account. Genome sequence sizes were measured as the number of unambiguous nucleotides in the assembly. A set of 18 commonly used palindromic type II restriction enzymes with variable nucleotide compositions was screened in each of the genome assemblies (Table 1). The number of cleavage sites present in each genome was obtained by counting the number of unambiguous matches for each recognition sequence pattern. Under optimal experimental conditions each cleavage site should produce two RAD tags, one in each direction from the restriction site. Therefore, we define the number of observed RAD tags in each genome assembly as twice the number of recognition sequence pattern matches.

### Recovery of restriction-site associated DNA tags

The number of cleavage sites in a genome is not the only factor that determines the number of RAD loci that can be recovered experimentally. The architecture of each genome, and in particular the number of repetitive elements and gene duplications, can significantly decrease the number of unambiguous loci obtained via alignment to a reference genome or *de novo* assembly. To quantify this contribution we assessed the proportion of restriction-site associated DNA tags that can potentially be recovered unambiguously after empirical sequencing. We performed *in silico* sequencing experiments for all genome assembly-restriction enzyme combinations. For each restriction site located in the genome assemblies, 100 base pairs up- and down-stream of the restriction site were extracted. This sequence read length is typical of sequencing experiments performed with current Hi-Seq platforms (Illumina Inc.). The resulting RAD tags were aligned back to their original genome assemblies using BOWTIE v0.12.7 (Langmead *et al.* 2009). Only reads that produced a unique best alignment were retained.

### Genome size estimation

To explore the potential for predicting genome sizes from restriction recognition-sequence frequency data, we modeled their relationship using data from the 434 genomes and 18 restriction enzymes through linear regression. Genome sizes and restriction recognition-sequence frequencies were log_10_ converted to handle the multiple orders of magnitude spanned within each variable. The non-parametric Spearman’s rank-order correlation coefficient (*ρ*) was calculated to measure the strength of association between genome sizes and restriction recognition-sequence frequencies. Simple linear models were fitted using least-squares estimation of *β* parameters with the *lm* function in R. The generalized simple linear model (1) used to predict genome size *y*, in units of base pairs, is defined as:

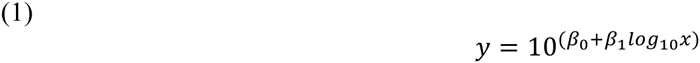

Where *x* is the number of restriction recognition-sequences in the genome, and *β*_0_ and *β*_1_ are the estimated parameters. Table 2 provides the estimated values of *β*_0_ and *β*_1_ for each restriction enzyme.

### Restriction recognition-sequence probability calculation

To test the hypothesis that compositional heterogeneity in eukaryotic genomes can determine the frequency of cleavage sites of each genome, we characterized the GC content, as well as the mononucleotide, dinucleotide and trinucleotide compositions of each genome and developed probability models to predict the expected frequency of recognition sequences for each restriction enzyme. GC content was calculated as the proportion of unambiguous nucleotides in the assembly that are either guanine or cytosine, assuming that the frequency of guanine is equal to the frequency of cytosine. Mononucleotide composition was determined as the frequency of each one of the four nucleotides. Dinucleotide and trinucleotide compositions were determined as the frequency of each one of the 16 or 64 possible nucleotide combinations, respectively.

Mononucleotide and GC content sequence models were used to estimate the probability of a particular recognition sequence assuming that each nucleotide is independent of the others and of its position on the recognition sequence. The GC content model assumes that the relative frequencies of guanine and cytosine in the genome sequence are equal. This model has only two parameters, the GC and AT frequencies. In the mononucleotide model (2) there are four parameters, one for each of the four possible nucleotides.

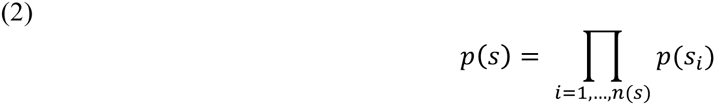

Here, *p(s_i_)* is the probability of nucleotide *s*_*i*_ at the position *i* of the recognition sequence. In the GC content model *p(s_i_)* can take the values of *f*_*G, C*_ or *f*_*A, T*_. In the mononucleotide model *p(s_i_)* can take the values of *f*_*A*_, *f*_*G*_, *f*_*C*_, or *f*_*T*_, where *f*_*X*_,is the frequency of a given mononucleotide (*X = A, G, C, or T*).

Dinucleotide and trinucleotide sequence models (3) were defined as first and second degree Markov chain transition probability models with 16 or 64 parameters, respectively (Karlin et al. 1992; Singh 2009). These models take into account the position of each nucleotide in the recognition sequence. Nucleotides along the recognition sequence are not independent from nucleotides in neighboring positions. The probability of a particular recognition sequence for these Markov chain models was calculated as:

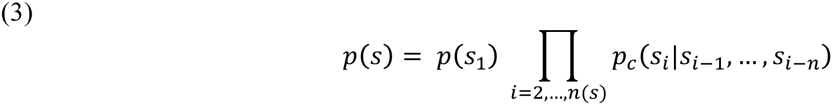

Where *p*_*S*1_ is the probability at the first position on the recognition sequence and *p*_*c*_ is the conditional probability of a subsequent nucleotide on the recognition sequence depending on the previous *n* nucleotides. In the dinucleotide sequence model *n* = 1 and in the trinucleotide sequence models *n* = 2.

Genomic resources are unavailable for most species. However, reduced representation datasets that capture a small fraction of a genome, such as RNA-seq or RAD-seq datasets, are more widely available. We investigated the potential use of these datasets to estimate genome composition parameters for our predictive models and calculate recognition-sequence probabilities for the selected set of 18 restriction enzymes. For this we selected a set of 27 species out of the 434 examined eukaryotic species with whole and draft genomes, which also have publically available transcriptome data (Supplementary Table 2). We also used the data from the *in silico* RAD sequencing experiments (described above) as reduced representation datasets for these species. We estimated genome composition parameters from transcriptome and RAD-seq datasets, and calculated recognition-sequence probabilities using the models.

### Expectations versus observations

To assess the effectiveness of the predictive recognition sequence models, we compared the number of observed restriction sites (frequency) in the genome assemblies with the expected predicted number according to each model, parameters estimated from whole and draft genome datasets. The expected number of restriction sites in a given genome was calculated as the product of the probability of a recognition sequence multiplied by the genome sequence size. To quantify the departures from expectation, we define a similarity index (*SI)* as *SI =* (*O–E*)*/E*, where *O* and *E* are the observed and expected number of restriction sites, respectively. If *SI =* 0, then *E = O.* If *SI <* 0, then *E > O,* and *vice versa*.

To measure the overall similarity between the restriction recognition-sequence probabilities calculated using known composition parameters from the genome and those calculated using estimated composition parameters from reduced-representation transcriptome and genome datasets, we calculated the mean squared error (MSE) per species.

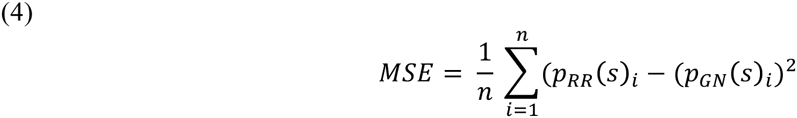

Where *p*_*RR*_(*s*)_*i*_ is the probability of a restriction recognition-sequence (of an enzyme *i*) calculated using composition parameters estimated from reduced-representation datasets, *p*_*GN*_(*s*)_*i*_ is probability of a restriction recognition-sequence calculated using known composition parameters from genome datasets. Each enzyme was assigned an arbitrary number from 1 to 18 (*n*). When MSE=0 the probabilities are identical. MSE value increases as similarity decreases.

### Location of recognition sequences in mammalian genomes

To evaluate the possibility that recognition-sequence frequency patterns inconsistent with evolutionary neutrality occurred in genomic areas subject to natural selection, we investigated the genomic locations of recognition sequences relative to well-annotated conserved-element genomic regions. We obtained DNA sequences of genomic elements (*sensu* Siepel *et al.* 2005; Miller *et al.* 2007b) strongly conserved across mammals from the human, dog and mouse genomes using the UCSC genome table browser (http://genome.ucsc.edu/cgi-bin/hgTables). We counted the number of occurrences of recognition sequences for each of the 18 restriction enzymes in these conserved genomic elements (observed) and compared them, using the similarity index (SI) described above, to the expected number of occurrences in a random genome sample of equal size (calculated as the relative frequency of recognition sequences in the whole genome [total number of recognition sequences/genome size in base pairs] multiplied by the size of each of the conserved-elements datasets in base pairs).

The analytical software pipeline here described (PredRAD), visualization scripts, and output database files are publicly available at https://github.com/phrh/PredRAD.

## SUPPLEMENTARY MATERIAL

Supplementary figures 1-9, and tables 1 and 2 are available as supplementary material.

## AUTHOR’S CONTRIBUTIONS

SH conceived and designed research. SH and PHR developed the software. SH analyzed the data. SH, TMS and PHR contributed computing equipment. SH wrote the paper with comments from PHR and TMS.

## ACKNOWLEDGEMENTS

This research was supported by the Office of Ocean Exploration and Research of the National Oceanic and Atmospheric Administration (NA09OAR4320129 to TS); the Division of Ocean Sciences of the National Science Foundation (OCE-1131620 to TS); the Astrobiology Science and Technology for Exploring Planets program of the National Aeronautics and Space Administration (NNX09AB76G to TS); and the Academic Programs Office (Ocean Ventures Fund to SH), the Ocean Exploration Institute (Fellowship support to TMS) and the Ocean Life Institute of the Woods Hole Oceanographic Institution (internal grant to TMS and SH). We thank Adam Reitzel and Ann Tarrant for helpful discussions. Ann Tarrant, Eleanor Bors, Annette Govindarajan, and Jill McDermott provided constructive comments on this manuscript.

## APPENDIX I

### Genomic composition patterns across the eukaryotic tree of life

The odds ratios proposed by Burge *et al.* (1992) were used to estimate compositional biases of dinucleotides (5) and trinucleotides (6) across genomes.

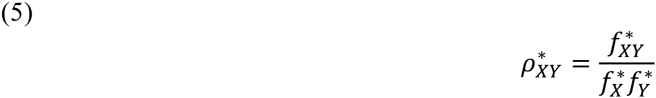

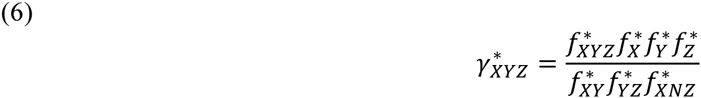

Where 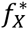 is the relative frequency of the mononucleotide *X,* 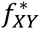 is the relative frequency of the dinucleotide *XY*, and 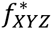 is the relative frequency of the trinucleotide *XYZ*. All frequencies take into account the antiparallel structure of double stranded DNA. *N* represents any mononucleotide. Both dinucleotides and trinucleotides are considered significantly underrepresented if the odds ratio is ≤ 0.78, significantly overrepresented if ≥ 1.23, and equal to expectation if =1 (Karlin et al. 1998).

Our surveys of whole and draft genome sequence assemblies indicate that there are significant compositional biases for most dinucleotides and trinucleotides across the eukaryotes. Many of these biases are significant only within individual species scattered throughout the eukaryotic tree of life. However, there are several particular dinuclotides and trinucleotides that show significant biases across the eukaryotic tree of life. The dinucleotides CG, GC, TA, and CA/TG, and the trinucleotides CTA/TAG, AAA/TTT, TAA/TTA, CCA/TGG show the most conspicuous bias patterns. Our observation that these biases are highly variable among broad taxonomic groups but generally similar within is congruent with findings from previous studies (e.g., Gentles & Karlin 2001). The most obvious biases across taxa are observed in the gnatostomate vertebrates; however, this is most likely due to rampant undersampling in most other groups of eukaryotes (vertebrate genome assemblies represent 21% of all the taxa in this study).

### Dinucleotide compositional biases

Dinucleotide odds ratios 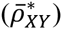 (Burge et al. 1992), a measurement of relative dinucleotide abundances given observed component frequencies used to explore genomic compositional biases, revealed significant compositional biases for all possible dinucleotides (Supplementary Figure 7). The dinucleotide compositional biases were highly variable among broad taxonomic groups (e.g., core eudicot plants) but generally similar within. Two dinucleotide complementary pairs, CG/GC and AT/TA, had highly dissimilar relative frequencies between the members of each pair. The largest biases were for CG, being significantly underrepresented in groups like core eudicot plants 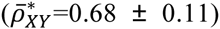, gnathostomate vertebrates 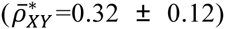, the Pucciniales rust fungi 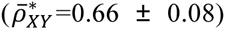, gastropod mollusks 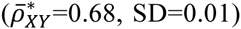, the Trebouxiophyceae green algae 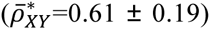 and the Saccharomycetales yeast 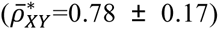. CG was significantly overrepresented in groups like the Apocrita insects 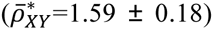. The complementary dinucleotide GC was not particularly underrepresented in any broad taxonomic group, but tended towards overrepresentation in ecdysozoan invertebrates 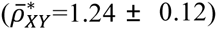, being significant in several arthropod and nematode species. Other taxa that showed significant overrepresentation of GC dinucleotides included the Trebouxiophyceae 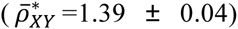 and microsporidia fungi 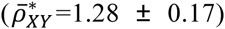 Relative abundances of the dinucleotide AT were within expectations for all eukaryotes, except for the fungus *Sporobolomyces roseus* 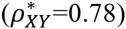 Contrastingly, the TA dinucleotide tended towards underrepresentation throughout the eukaryotes 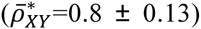, except in a few hypocreomycetid fungal species, for which it was significantly underrepresented. The TA dinucleotide was significantly underrepresented in trypanosomatids 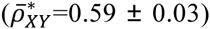, choanoflagellids 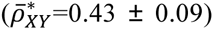, chlorophytes 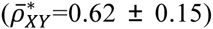, stramenopiles 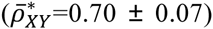, and marginally underrepresented in most euteleostei fish 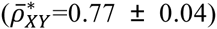, archosaurs 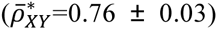 and the Basidiomycota 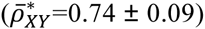, among others.

The remaining dinucleotides had identical relative frequencies between the members of each complementary pair. The dinucleotide pair GG/CC was marginally underrepresented in most eukaryotes 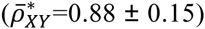. In the Sarcopterygii vertebrates 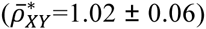 and embryophyte plants 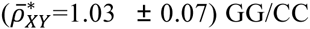 GG/CC relative frequencies closely conformed to expectation, whereby GG/CC was significantly overrepresented in handful of isolated ecdysozoan, microsporidia and alveolate species, and significantly underrepresented in chlorophytes 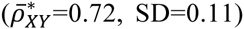, oomycetes 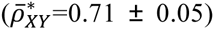, and in several species of the Basidiomycota and the Dothideomycetes. Only the choanoflagellate *Salpingoeca* and the green alga *Asterochloris* presented a marginally significant bias for the dinucleotide pair AA/TT 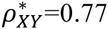. Similarly, *Salpingoeca* was the only taxon to show a significant bias for AC/GT 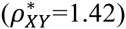). Dinucleotide pair CA/TG was among the pairs with largest biases. Significant overrepresentation of CA/TG was found in several groups with large CG underrepresentation such as gnathostomates 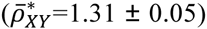, gastropods 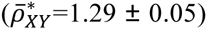, the Pucciniales 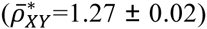, the Trebouxiophyceae 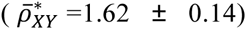, as well as several species of core eudicots and the Saccharomycetales. Other groups with significant CA/TG overrepresentation include onchocercid nematodes 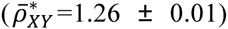, the Ustilaginomycotina fungi 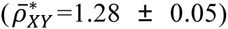, trypanosomatids 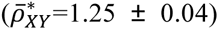, and amoebozoans 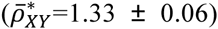. Overrepresentation biases for the AG/CT dinucleotide pair were only present in amniotes 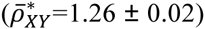, the Sporidiobolales fungi 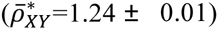, and oxytrichid alveolates 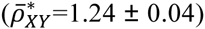, and other isolated species. Most of these taxa also had large CG underrepresentation. Lastly, most eukaryotes had GA/TC relative frequencies that conformed to expectations, except for few scattered species and small groups such as the Microbotryomycetes fungi 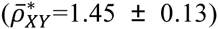, the Mamiellales green algae 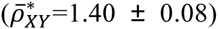, and the Eimeriorina alveolates 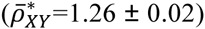.

Biases in most of these dinucleotides are likely linked to important biological processes. Notably the underrepresented dinucleotide CG is a widely known target for methylation related to transcriptional regulation (Bird 1980) and retrotransposon inactivation (Yoder et al. 1997) in vertebrates and eudicots. The corresponding overrepresentation of AG/CT fits the classic model of “methylation-deamination-mutation” by which a methylated cytosine in the CG pair tends to deaminate when unpaired and mutate into a thymidine with a corresponding CA complement. Interestingly CG and GC dinucleotides are significantly overrepresented in several groups of apocritic insects, as well as in some fungi and single-cell eukaryotes. CG is not a primary target for methylation in *Drosophila* (Lyko et al. 2000), instead CT, and in lesser degree CA and CC, are methylated in higher proportion. None of these dinucleotide pairs is significantly underrepresented in apocritic insects. The widespread TA underrepresentation has been traditionally attributed to stop codon biases, thermodynamic instability and susceptibility of UA to cleavage by RNAses in RNA transcripts (Beutler et al. 1989).

### Trinucleotide compositional biases

Trinucleotide odds ratios 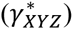 (Burge et al. 1992) are another important measurement used to explore genomic compositional biases. Among the examined taxa, these ratios revealed compositional biases for most possible trinucleotides (Supplementary Figure 8). However, most of these biases were only significant in scattered individual species (Supplementary Figure 9). Among the trinucleotide pairs with significant underrepresentation, CTA/TAG and CGA/TCG showed the most definite broad taxonomic patterns. CTA/TAG was significantly underrepresented in most taxa, except for groups like commelinid plants (monocots) 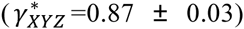, most core eudicots 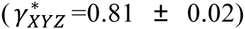, eleutherozoans 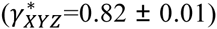, molluscs 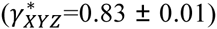, and gnathostomates 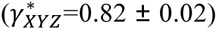 – exclusive of the chimaera *Callorhinchus milii*. Contrastingly, the trinucleotide CGA/TCG was only significantly underrepresented in most tetrapod vertebrates 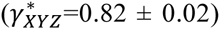[g3]=0.82 ± 0.02) – exclusive of muroid rodents, bovid ruminants and the Afrotheria – a group containing aarvdvarks, hyraxes, and elephants.

The largest and more widespread overrepresentation biases were for the trinucleotide pair AAA/TTT, being significant in most eukaryotes, except for the majority of the Dikarya fungi 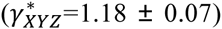. The trinucleotide pairs TAA/TTA and AAT/ATT were significantly overrepresented in many metazoan taxa, particularly in the Neopterygii vertebrates 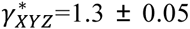, and 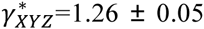. AAG/CTT was significantly overrepresented in the Bacillariophyta diatoms 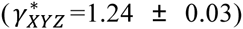, oomycetes 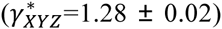, and the Saccharomycetales 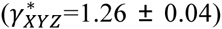. Lastly, CCA/TTG was significantly overrepresented in several tetrapod groups, including the Laurasiatheria – exclusive of the Chiroptera – 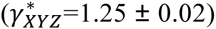 and Hominoidea 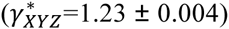.

The biases in CTA/TAG have been widely attributed to the stop codon nature of UAG. However, the trinucleotides corresponding to the other stop codons (Burge et al. 1992), UAA and UGA, are overrepresented or not biased across eukaryotes. The reasons behind other cases of trinucleotide biases are less understood.

